# A rapid HPLC-based method to determine NAD(P)(H) and NMN redox cofactor concentrations and ratios in microbes

**DOI:** 10.64898/2026.07.24.740514

**Authors:** Noor E. van Wijk, Elisabeth C.M. van der Heijden, Javier M. Hernández-Sancho, Daniel C. Volke, Pablo I. Nikel, Johan H. van Heerden, Frank J. Bruggeman, Nico J. Claassens, Ruud A. Weusthuis, Markus M.M. Bisschops

## Abstract

Redox cofactors are a key part of cellular physiology as they are involved in most metabolic pathways, and their ratios are linked to cellular robustness. However, measuring their levels in cells remains a challenge. Here, we describe a novel method to rapidly measure NAD(H), NADP(H) and nicotinamide mononucleotide (NMN) levels and their oxidized/reduced ratios using an HPLC connected to a fluorescence detector. By extensively characterizing this method and benchmarking it against the classical iodonitrotetrazolium (INT) assay, we show that this method results in accurate and reproducible measurements of NAD^+^, NADP^+^ and NMN levels in bacteria. We further demonstrate that this method can be used to determine intracellular NADH and NADPH concentrations and ratios of nicotinamide nucleotide cofactors in engineered *Escherichia coli* strains, as well as other bacterial species such as *Pseudomonas putida*.

## Introduction

Redox cofactors play essential roles in electron transfer and metabolic regulation in all cells, with NAD(H) and NADP(H) being the most abundant ones. Ratios of NADH/NAD^+^ and NADPH/NADP^+^ determine the direction of many reactions and thereby the driving force of pathways ^1–4^. In addition, NAD functions as substrate for a number of non-oxidoreductase enzymes and consequently plays an important role in DNA repair, NAD-capping of RNA, regulation of gene expression, stress response and cell survival ^5,6^. Therefore, altered redox ratios and concentrations in cells can have a large impact on cell physiology. For example, changes in NAD homeostasis have been linked to several human diseases, as well as aging ^7^. In microbial biotechnology, non-optimal cofactor levels and ratios can result in an inefficient flux through native and engineered pathways and thus lower production rates and yields ^8–10^. This makes the levels and ratios important targets for understanding cellular physiology, as well as metabolic engineering. In addition, cofactor levels and ratios also respond to, for example, oxidative stress ^11^, and nutrient limitations^12^. Thus, it is important to quantify these cofactors to gain a good understanding on cell physiology which is required for many fields working with cells.

Compared to the vast amount of metabolic engineering studies that involve redox reactions, the number of studies that actually measure redox cofactors’ levels or ratios is relatively limited ^13^. This is, at least in part, because quantifying redox cofactors is challenging due to their instability, quick enzymatic turnover, and low concentrations ^14^. Therefore, a method is needed that is easy, precise, compatible with quick quenching methods, applicable to cells of different microbial and other species, and affordable to make it accessible for measuring cofactors in many samples and laboratories.

Different methods have been described to determine cofactor levels or ratios. In the past decade, genetically encoded biosensors have been developed for NAD^+^, NADH, NADP^+^ and NADPH or their ratios NADH/NAD^+^, NADPH/NADP^+ 15–23^. Although relatively fast, inexpensive, and compatible with high-throughput applications, these biosensors have limitations, including cross-reactivity with related metabolites, restricted dynamic range, sensitivity to variables such as pH and ionic composition, reliance on molecular oxygen for chromophore maturation, and limited capacity for absolute quantification of intracellular cofactor levels. In addition, these biosensors require genetic engineering, which limits their applicability to genetically amenable cells, but may also impact cell performance and require thorough calibration.

To more exactly quantify cofactor levels, several colorimetric or fluorometric enzyme assays are available, including the Bernofsky and Swan method, fluorometric assays based on resazurin or commercial colorimetric/fluorometric kits ^24–26^. However, these assays also require extensive fine tuning and are quite labor-intensive. In addition, each assay only allows one redox-cofactor (pair) to be quantified, which means that to measure both NAD^+^/NADH and NADP^+^/NADPH at least two assays need to be performed.

To simultaneously measure all nicotinamide redox cofactors, LC/MS-based methods are often used ^27,28^. LC/MS cannot only quantify redox cofactors, but also many other metabolites. Therefore, LC/MS methods are often optimized for metabolomics, rather than measuring specifically redox cofactors only. Since NAD(H) and NADP(H) are quite similar, they are difficult to separate, and dedicated protocols have been developed to specifically quantify these and other metabolites in the NAD^+^ metabolome ^14,27,29,30^. Major drawbacks, however, are that these methods require expensive equipment, specialized personnel, and time for analysis of data. These disadvantages limit the throughput of this method for analysis of cofactor levels in microorganisms for metabolic engineering purposes.

A lesser used method is based on the reaction that turns NAD(P)^+^ and certain precursors into highly fluorescent 1,6-naphthyridine derivatives after condensation with a ketone ^31–34^. Following this approach, NAD(P)^+^ and precursors have been measured in enzyme assays and in urine, plasma and mouse tissues for clinical purposes ^31–34^. In addition, the concept has also been used to measure overproduction of NMN, an intermediate in NAD biosynthesis, in microbial cultures^35–37^. However, most of these studies used plate readers to measure the fluorescence, which means that results reflect changes in combined NAD(P)^+^ and NMN pools. To distinguish between different molecules, they can be separated by HPLC and measured with a fluorescent detector ^31–33^. Although this strategy could be used to measure NAD^+^ and NADP^+^, it has never been adapted to measure their reduced counterparts.

Here, we describe a method that quantifies NAD^+^, NADP^+^, NADH, NADPH, and NMN using HPLC with a fluorescence detector. With this method, we aim to make measuring intracellular redox cofactor levels and ratios more accessible, such that it can be widely used within nicotinamide redox cofactor research, such as in cellular physiology studies and metabolic engineering of microorganisms. We determined the cofactor levels in exponentially growing *Escherichia coli* cultures on different carbon sources, glucose, glycerol or acetate, and compared our results with results obtained using an enzymatic cycling assay. To show that this method is applicable to measure cofactors for microbial physiology and metabolic engineering purposes, we also used it to measure cofactor levels in engineered *E. coli* and *Pseudomonas putida* strains.

## Methods

### *E. coli* strain and plasmid construction

*E. coli* strains and plasmids used in this study are listed in Table S1. Primers used in this study are listed in Table S2.

The *pncC* gene was deleted by amplifying the kanamycin resistance cassette flanked by FRT sites and homology region from the genome of the JW2670-1 strain from the KEIO collection^38^ using the primers KO_PncC_R and KO_PncC_F and Phire polymerase (Thermofisher). Deletion of the gene was done as described in Jensen et al. (2015) ^39^.

The codon harmonized^40^ *nadV* gene from *Francisella tularensis* (FtNadV) was ordered as a gblock at Twist Bioscience (see supplementary information for sequence). Plasmids were built using the Sevab3.1 plasmid system^41^. For this, FtnadV was first cloned into a repository vector pSb1C3, for which the backbone was amplified with primers EV and PV. The gene was amplified with primers OP1_BBaB0034_FtNadV and SEVA_FtNadV_Rv. The pSEVAb32 backbone was amplified with SEVA_EvBBa_J23100 and PV. Q5 polymerase was used for amplification of DNA fragments for plasmid construction. Golden Gate assembly of fragments was done using 1 μL BsaI, 0.5 μL T4 DNA ligase and a thermocycling program (37 °C, 20 min; (16 °C, 4 min; 37 °C, 3 min) x 30; 50 °C, 10 min; 80 °C, 10 min).

*E. coli* DH5α λpir was transformed using heat shock for plasmid propagation. Aliquots of 50 μL of chemically competent *E. coli* DH5α λpir were mixed with 5 μL of golden gate reaction. The cells were incubated for 20 minutes on ice, followed by a heat shock at 42 °C and incubation on ice for 5 minutes. The cells were recovered in LB for 1 hour at 37 °C. Finally, the mixture was plated on LB agar plates with appropriate antibiotics.

### *E. coli* cultivation

*E. coli* strains were cultured on Lysogeny Broth (LB) at 37 °C, 250 rpm overnight. Cultures for analysis grew on minimal M9 medium with different carbon sources. M9 contained per liter: Difco^Tm^ M9 salts (BD Biosciences):12.4 g Na_2_HPO_4_•7H_2_O, 3.0 g KH_2_PO_4_, 1.0 g NH_4_Cl, 0.5 g NaCl; Trace elements: 8.30 mg FeCl_3_ •6H_2_O, 0.84 mg ZnCl_2_, 0.13 mg CuCl_2_•2H_2_O, 0.10 mg CoCl_2_•6H_2_O, 0.10 mg H_3_BO_3_, 0.016 mg MnCl_2_•4H_2_O; 0.2 mM MgSO_4_; 2 mL 1 M MgSO_4_ and 100 µL 1 M CaCl_2_. Carbon sources were 20 mM glucose or 40 mM glycerol or 60 mM acetate. When required, 30 µg ml^-1^ chloramphenicol or 50 µg ml^-1^ kanamycin was added.

Cultures were grown in 50 mL culture volume in 250-mL or 500-mL Erlenmeyer flasks. Growth was monitored by measuring optical density at 600 nm with a Hach DR6000 spectrophotometer or an Amersham Ultrospec 2100 pro spectrophotometer. The conversion factor to convert OD_600nm_ to gDCW was determined by measuring dry cell weight. This conversion factor was found to be 0.46 OD_600nm_ gDCW^-1^ (Hach DR6000) or 0.36 OD_600nm_ gDCW^-1^ (Amersham Ultrospec 2100 pro).

### Measuring fluorescent derivatives of the cofactors with HPLC-FLD and platereader

#### Materials for standard solutions

For preparation of standard solutions, the following chemicals were used: β-nicotinamide adenine dinucleotide sodium salt (Sigma-Aldrich) (NAD^+^), β-nicotinamide adenine dihydrate dinucleotide disodium salt (NADH) (Sigma-Aldrich), NADP disodium salt (Millipore) (NADP^+^), NADPH tetrasodium salt (Millipore), β-nicotinamide mononucleotide (Sigma-Aldrich) (NMN), and 1-methylnicotinamide chloride (Sigma-Aldrich) (CH3-NAM). NAD^+^ and NADP^+^ were dissolved in MQ to a final concentration of 1 mM. NMN and CH3-NAM were dissolved in MQ to a final concentration of 20 mM. NADPH and NADH were dissolved in 0.1 M Tris buffer (pH 8) to a final concentration of 1 mM. CH3-NAM was stored at –20 °C and all other standard stocks were stored at –80 °C.

### Plate reader assays

Initial tests were performed using a plate reader to test the condensation reaction and quenching for NAD^+^, NADH, NADP^+^, NADPH, and NMN. Standards with final concentrations of 1 μM, 2.5 μM, 5 μM and 10 μM were prepared in MQ. To test oxidation of NADH and NADPH with PMS, 60 μL of 2 mM PMS was added to 1 mL of standard and left in the dark for 10 minutes. To test degradation in acidic or alkaline conditions, 53 μL of 4 M KOH or HCl was added to 1 mL of standard. Standards were heated for 5 minutes at 100 °C in a heating block and left to cool for 10 minutes at room temperature. pH was neutralized by adding 53 μL of 4 M HCl or KOH and 110 μL 2 M HEPES pH 7. Finally, to these standards also 60 μL of 2 mM PMS was added and incubated in the dark.

Fluorescent derivatization was done in 96-well plates as described by Marinescu et al. (2018) ^35^. Per well, 74 μL of standard was pipetted, 23.3 μL of 20% acetophenone in DMSO was added and 23.3 μL of 2 M KOH. The plate was kept at 4 °C for 2 minutes. Afterwards, 125 μL of 88% formic acid was added. The plate was incubated at 37 °C for 10 minutes. Fluorescence was measured directly after incubation in a BioTeck Synergy H1 (Agilent) plate reader at an excitation wavelength of 382 ± 20 nm and emission wavelength of 445 ± 20 nm.

#### Quenching, extraction and derivatization

Samples were taken during the exponential growth phase at OD_600nm_ 0.7 – 0.8 after at least four doublings. Standards were diluted in M9 with 20 mM glucose or 40 mM glycerol or 60 mM acetate, depending on the experiment. Samples of 1 mL of culture and standards were transferred into a 1.5-mL Eppendorf tube containing 53 μL of 4 M KOH or 4 M HCl and immediately vortexed. At the same time as processing the samples, standards for the standard curve were processed, to ensure the exact same handling and processing times as the samples.

Samples and standards were heated for 5 minutes at 100 °C in a heating block. Afterwards, they were left at room temperature to cool for 10 minutes. Samples were neutralized with 53 μL of 4 M KOH or 4 M HCl and 110 μL 2 M HEPES pH 7. To oxidize the reduced redox cofactors, 60 μL of 2 mM PMS was added and incubated for 10 minutes in the dark. To all samples, 3.2 μL of 1 mM CH3-NAM was added as internal standard. Samples were centrifuged for 10 minutes at 21130 rcf at 4 °C. 148 μL of supernatant was transferred to new Eppendorf tubes for the fluorescent derivatization. 56.6 μL of 20% acetophenone in DMSO and 56.6 μL of 2 M KOH were added. The tubes were incubated at 4 °C for 2 minutes. Next, 250 μL of 88% formic acid was added and incubated at 37 °C for 10 minutes. Derivatized samples were stored at –20 °C and analyzed within one week.

#### HPLC-FLD method

The HPLC protocol was adapted from Mori et al. (2014). Samples were analyzed with an Agilent Technologies 1290 Infinity system with a fluorescent detector. The guard column and column used were a Security guard C18 (4 mm x 3 mM, Phenomenex) and a Luna 5 μm C18 column (100 A, 150 x 4.6 mm), respectively. 20 μL of sample was injected into the HPLC. The fluorescent detector was set at an excitation wavelength of 380 nm and an emission wavelength of 440 nm. A gradient was used to eluate samples consisting of buffer A for 6 minutes at 100%, 5 minutes changing up to 100% buffer B, followed by another 6 minutes of 100% buffer B, followed by re-equilibration of buffer A for 6 minutes. Buffer A contained 100 mM KH_2_PO_4_ and 10% acetonitrile at a pH of 2.1. Buffer B contained 100 mM KH_2_PO_4_ and 40% acetonitrile at a pH of 2.1. The flow rate was 1.5 mL minute^-1^, and the column temperature was 25 °C.

### Method verification with HPLC-FLD

To verify the protocol, spiking experiments were performed. Cell culture samples of 990 μL were added to 53 μL of 4 M KOH or 4 M HCL and rapidly spiked with 10 μL of standards with known concentrations. Samples were further processed by the protocol described above.

### Data processing and visualization – HPLC-FLD

Data analysis was performed in Microsoft Excel and in GraphPad prism (version 10.6.1). Graphs were made in GraphPad prism. Peaks detected in the fluorescence detector were integrated with the LabSolutions software from Shimadzu. Standard curves were drawn by calculating the area of the peaks with standards containing 0.01 – 5 μM with nonlinear regression by fitting a straight line in GraphPad prism.

Limit of quantification (LOQ) and limit of detection (LOD) were calculated based on the signal-to-noise ratio S/N. The LOQ was determined as 3.3 = S/N and for LOD this was 10 = S/N using the standard curve made from standards diluted in M9 with 20 mM glucose. The average noise in the standards was determined in the Labsolution software (Shimadzu). Next, we calculated the signal (S) that was required for the LOD and LOQ. Based on a standard curve determined from the peak heights with linear regression, LOD and LOQ were calculated:

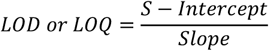

Here, S is the signal that was required for the 3.3 = S/N and 10 =S/N.

To evaluate the preciseness of the method, samples were spiked. Fluorescence of the spiked metabolite in the samples were calculated by:

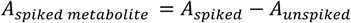

In which the area of the peak from the spiked sample (A_spiked_) was subtracted by the area of the peak from the un-spiked sample (A_unspiked_). The relative recovery of each spiked concentration was calculated by:

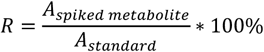

Here, A_standard_ is the peak area of the target cofactor measured in the standards that had the same concentration as the amount of cofactor spiked into the sample. Finally, coefficient of variation in % was determined to assess the preciseness of the method:

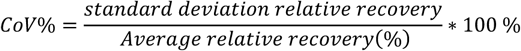

### Measuring cofactors with an INT cycling assay

The INT cycling assay protocol was adapted from the MTT cycling assay by Bernovsky and Swan (1970)^25^. The assay can be used to biochemically quantify cellular NAD^+^ and NADH concentrations and ratios, by making use of the fact that NADH is degraded in dilute acid and NAD^+^ in dilute alkaline solutions. In this assay, reduction of iodonitrotetrazolium chloride (INT) is measured, which is linearly proportional to the concentration of NAD(H). Here, we adapted the original protocol to a standard addition method, which is ideal for determining concentrations of compounds in complex mixtures, such as cell samples.

#### Quenching and extraction

Samples were taken in the exponential phase at an OD_600nm_ of 0.7-0.8 after at least four doublings. Samples were taken by transferring 10 mL of culture into 50-mL Greiner tubes with 0.532 mL of 4 M KOH or 4 M HCl. Subsequently, 1 mL aliquots were added to 2-mL Eppendorf tubes containing 10 μL of 0, 0.2, 0.15, 0.10 or 0.05 mM of NAD^+^ or NADH respectively in water. Additionally, 2 mL of culture was filtered using a 0.22 µm filter, after which 950 μL of the filtered spent medium was added to 50 μL 4 M KOH or 4 M HCl in 2 mL Eppendorf tubes to use as reagent blank (RB). All samples were transferred to a 100 °C heating block for 5 minutes to ensure complete breakdown of NADH or NAD^+^ in samples containing HCl or KOH, respectively. After the samples were spun down at 3100 rcf and 4 °C for 10 minutes in a Heraeus Fresco 17 tabletop centrifuge to remove cellular debris. The supernatant was neutralized by adding 1:1 (v/v) 1 M HEPES buffer pH 7.5 or pH 7.0 for HCl and KOH samples respectively. The pH of all samples was checked and adjusted if needed to ensure identical pH – around 6.5 for HCl and around 7.5 for KOH - between samples.

#### Preparation of the INT reaction mix and plate-reader measurement

On the day of analysis, 10 mL reaction mix per plate was freshly prepared in subdued light, containing 1 mL 500 mM Tris-HCl pH 7.5, 1 mL 40 mM EDTA, 2 mL color reagent, 1.2 mL ADH 500 mL^-1^ in 1 M bicine/NaOH pH 8.0 and 4.8 mL MQ. The color reagent, consisting of 1.63 mM phenazine methosulfate (PMS), 3.96 mM iodonitrotetrazolium chloride (INT), 2% (v/v) Triton X-100 and 6 M ethanol was prepared in batch and stored at 4 °C, protected from light. To fully dissolve all components, a sonication bath was used.

Extracted samples were vortexed, after which 10 μL aliquots were added to a 96 wells plate in 6 technical replicates of each concentration. Subsequently, a 90 μL reaction mix was added per well in subdued light conditions and instantly transferred to a BMG Labtech SPECTROstar Nano microplate reader. Reduction of INT was measured at 490 nm, 20 flashes per measurement, 37 °C, without shaking, and a settling time of 0.2 seconds. Measurements were taken every 50 seconds for approximately 30 minutes or until the signal stabilized. NAD(H) levels are determined based on the linear relation between slope and concentration.

### Data processing and visualization – INT assay

Data analysis was performed in Microsoft Excel and Python (version 3.10.14)

Mean replicates of slopes in raw data (OD_490_/time) were calculated per sample by linear regression. These slopes were plotted against the concentration of standard added to the sample, which again resulted in a linear relation termed the standard addition plot. Concentrations of NAD(H) could be determined from the x-intercept of the latter plot.

The limit of quantification and limit of detection were calculated using the equations below, with IUPAC-recommended multipliers (3.3 and 10 respectively), as common for standard addition methods with heteroscedasticity in noise.

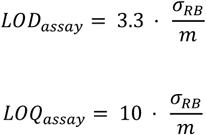

Where m is the slope of the standard addition plot, and σ_RB_ is the standard deviation of the slopes of the RB replicates. These values were converted to biologically relevant numbers

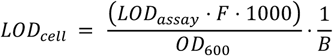

Where F is a conversion factor based on the dilution of cell samples (10/950) and B is a biomass constant of 0.36 gDCW L^-1^ per OD600 unit. For this assay the OD600 value of the cell samples was 0.7. As the noise in the RB replicates varied per carbon source used, this was repeated for every condition.

### Pseudomonas putida experiments

*P. putida* strains and plasmids used in this study are listed in Table S1.

Gene deletions were carried out via homologous recombination following an established protocol ^42^. For the I-SceI mediated recombination, the corresponding suicide pSNW2-derivative plasmid was delivered into the cells by electroporating around 300 ng of plasmid DNA into 80 µL of freshly prepared electrocompetent *P. putida* cells, which were prepared by washing the biomass from 10 mL of an overnight culture with 1 mL of 300 mM sucrose four times. Electroporation was performed with a Gene Pulser XCell (Bio-Rad) set to 2.5 kV, 25 µF capacitance, and 200 Ω resistance in a 2-mm gap cuvette. The co-integrants were subsequently transformed with pQURE6·H ^42^. Cells were recovered in 1 mL of LB medium supplemented with 2 mM of 3-methylbenzoate (3-mBz) for at least 2 h at 30 °C and plated onto LB medium agar containing gentamycin and 3 mM 3-mBz to induce both plasmid replication and I-SceI expression. Gene deletions were identified by colony PCR and verified by Sanger DNA sequencing. Finally, pQURE6·H was cured from the resolved co-integrant by omitting 3-mBz in subsequent passages of the culture.

*P. putida* KT2440, and the derived strain with the *pncC* deletion and harboring the plasmid pP4S·*nadV* were precultured in minimal salt medium (MSM) supplemented with 20 mM glucose ^43^. Afterwards, precultures were used to inoculate 50 mL of MSM medium supplemented with 20 mM glucose, 1 mM nicotinamide and 5 mM rhamnose in 500 mL Erlenmeyer flasks at an OD_600nm_ of 0.05. Growth was monitored by measuring optical density at 600 nm with a Hach DR6000 spectrophotometer and the cultures were harvested in the exponential growth phase (OD_600nm_ values between 0.4-0.8). 25 mL of culture were used to perform the INT cycling assay. The other 25 mL were pelleted and stored at −20 °C.

Pellets were resuspended in 1 mL of 1% (v/v) formic acid and cooled in ice water. Samples were transferred to a cryotube with 0.3 g of acid washed crystal beads (425-600 μm, Sigma-Aldrich). Cells were lysed during 3 cycles of 20 s at 6,500 rpm and 15 s of rest in a bead beater Precellys 24 Dual (Bertin Technologies SAS, Montigny Le Bretonneux, France). After lysis, the samples were immediately placed in ice water for 5 min and then centrifuged for 10 min at 4 °C and 17,000 *g*. The supernatant was stored at – 20 °C. 74 µL of the lysate was thawed on ice water and transferred to a new Eppendorf tube. After addition of 28 µL of 20% acetophenone in DMSO and 28 µL of 2 M KOH, the tubes were vortexed for 3 s and kept on ice for 2 min. This was immediately followed by addition of 125 µL of 88% (v/v) formic acid. The sample was vortexed for 3 s and incubated for 10 min at 37 °C. Subsequently, the samples were filtered using 0.22 µm syringe filters (Labsolute®, Th.Geyer, Renningen, Germany). HPLC spectrofluorometric analysis was performed using a sPFP column and 10 mM ammonium formate (eluent A) and acetonitrile (eluent B). The elution gradient started with 5% of eluent B at 0 min, increasing to 60% at 5 min and 90% at 5.5 min. The gradient allowed for baseline separation of NMN. Metabolites were detected using a fluorescence detector at 310 nm excitation and 430 nm emission wavelengths. Independent calibration curves were performed for each compound (NAD^+^ and NMN). The intracellular concentrations were calculated assuming cell dry weight (0.48 g L^-1^ OD_600nm_^-1^) ^44^.

## Results

### Reduced and oxidized cofactor pools can be efficiently separated by heating in either strong acid or base

The reaction to derivatize NAD(P)^+^ and precursors to highly fluorescent 1,6-naphthyridine derivatives has been optimized in multiple studies ^33,45^. So far, only fluorescent derivatization of the oxidized cofactor has been reported and not the reduced counterparts NADH and NADPH. Therefore, we tested whether NADH and NADPH could also be derivatized to become fluorescent. This was not the case under standard conditions (Figure 1A). To increase response in fluorescence, we added the strong oxidizer phenazine methosulfate (PMS) ^25,46^ and tested if it could completely oxidize NADH and NADPH to NAD^+^ and NADP^+^, respectively. Indeed, NADH and NADPH standards became equally fluorescent as NAD^+^ and NADP^+^ after the addition of PMS (Figure 1B).

**Figure 1.**
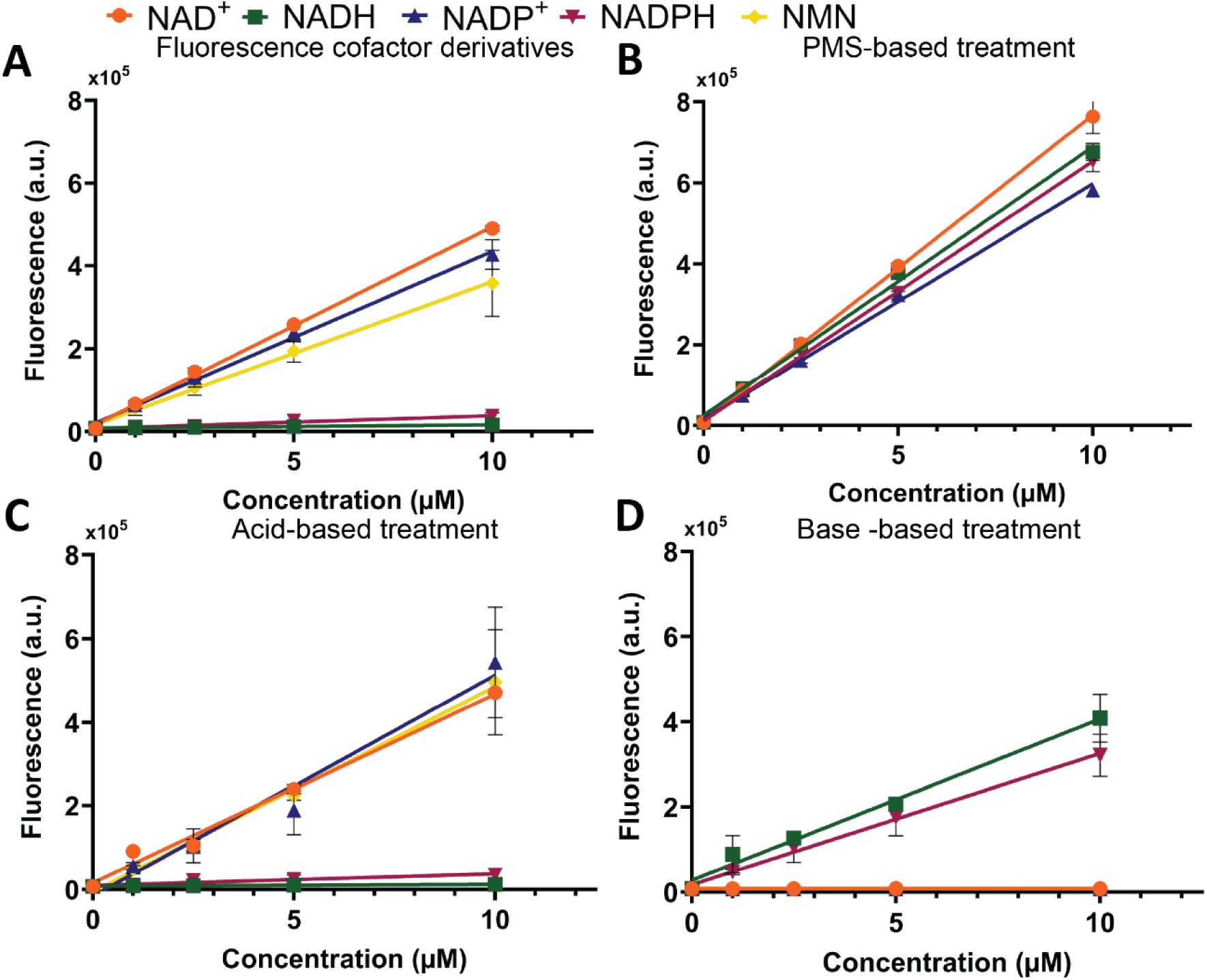
A) Fluorescence of NAD(P)(H) and NMN derivatives when dissolved in MQ B) Fluorescence of NAD(P)(H) derivatives after addition of PMS. C) Fluorescence of NAD(P)(H) derivatives after heating standards in 0.2 M HCl, neutralizing pH and addition of PMS. D) Fluorescence of NAD(P)(H) derivatives after heating standards in 0.2 M KOH, neutralizing pH and addition of PMS. Mean values and standard deviations, shown as error bars, were calculated from duplicate samples.

Oxidized cofactors are stable in low pH solutions, while the reduced cofactors are more stable in alkaline conditions ^47,48^. We tested if we could separate the oxidized and reduced cofactor pool by heating the standards in either acid or base. After this treatment, PMS was added to the samples prior to derivatization. No to negligible fluorescence was detected after NADH and NADPH were heated in acid (Figure 1C). Their degradation products did not turn fluorescent after derivatization, while fluorescence was detected for their oxidized counterparts. These oxidized molecules were degraded in alkaline conditions: NAD^+^ and NADP^+^ did not show any fluorescence after alkali treatment (Figure 1D). The reduced and oxidized cofactor pools could thus be separated by heating at either a high or low pH.

### Characterization and validation of the HPLC-FLD method

Next, we further developed the method to be able to simultaneously detect the different cofactors by separating molecules using HPLC and measuring their fluorescence with an FLD (Figure 2A). The different standards resulted in separate peaks (Figure 2 B, C). To check whether retention times and derivatization were constant between samples and standards, as well as between runs, methylnicotinamide (CH_3_-NAM) was used as an internal standard. To determine the concentration of the cofactors, the area under the highest peak for each metabolite was used (Figure 2 B, C).

**Figure 2.**
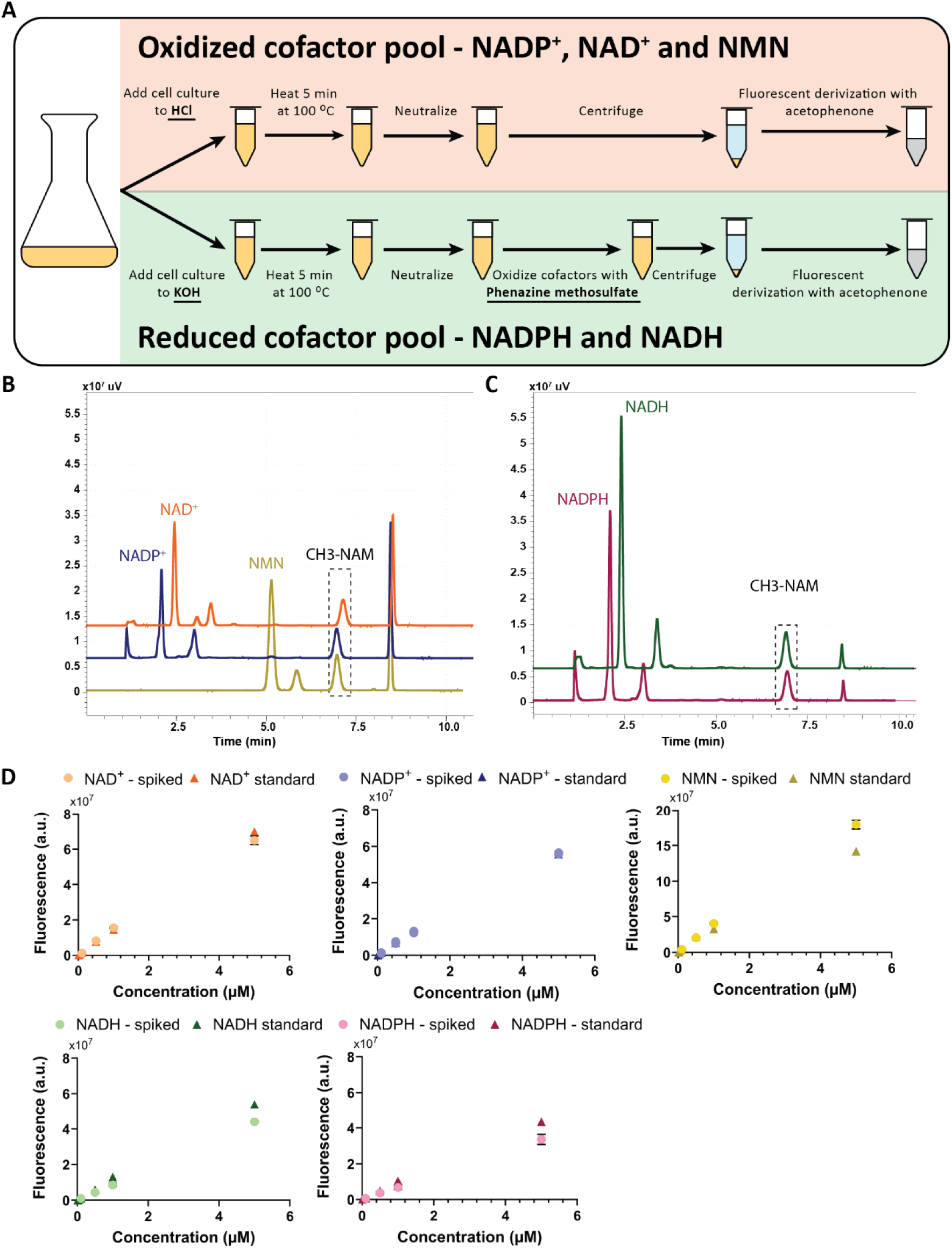
HPLC-FLD method characterization. A) workflow of protocol. B) Overlay of chromatograms of oxidized standards of 10 μM NAD^+^, NADP^+^ and NMN. C) Overlay of chromatograms of reduced standards of 10 μM NADH and NADPH. Standards contained methyl-nicotinamide as internal standard. D) Known concentrations of redox cofactors and NMN spiked to *E. coli* cultures (circles) compared to a standard curves prepared in M9 + 20 mM glucose medium measured as single replicate (triangles). For spiked samples, mean values and standard deviations, shown as error bars, were calculated from triplicate samples.

The robustness and preciseness of the protocol were evaluated based on the standard curves with samples ranging from 0.01 μM to 10 μM. For each metabolite, duplicate standard curves were prepared and analyzed over 3 different days (Figure S1). Minor variations were seen between standard curves for the same molecules, which might be due to small differences in mobile phase composition or fluorescent derivatization reaction times. Fluorescence intensity of the metabolites depends on the pH, so slight differences in pH of the mobile phase can impact the fluorescence intensity of the standards ^33^. Limit of detection (LOD) and limit of quantification (LOQ) were calculated using the S/N method with S/N=3.3 for LOD or S/N=10 for LOQ. Following this method, the LOD was around 0.01 μM and the LOQ around 0.05 μM for all cofactors(Table S3). For standard curves between 0 and 10 μM, linear regression resulted in R^2^ values between 0.96-1 (Figure S1).

To further validate the preciseness of this method in biologically relevant sample conditions, samples from *E. coli* cultures grown on minimal media with glucose and quenched in either HCl or KOH were spiked with known concentrations of the different cofactors. We clearly saw an increase in fluorescence in the samples that matched the fluorescence levels from the standards (Figure 2D). The average recovery for NAD^+^ and NADP^+^ is high (>90%) for the higher concentrations. Only when samples were spiked with 0.1 μM NAD(P)^+^ the measurements were not consistent resulting in a high coefficient of variation (CoV) (Table S4). For NMN, we saw similar results; however, we saw an overestimation of the highest spiked concentration.

The recovery of NADH and NADPH was much lower (∼70–80%) with a higher CoV for all concentrations compared to NAD^+^ or NADP^+^. A putative explanation is that NADH and NADPH could be degraded faster in a different matrix ^49^. To check the influence of medium composition, standard curves were prepared in M9 medium without glucose, with 20 mM glucose and in filtered medium obtained from exponentially growing *E. coli* cultures. The presence of glucose clearly led to lower fluorescence intensities compared to standards dissolved in M9 without glucose (Figure S2). Since the slope differed only 10% between spent medium and medium with 20 mM glucose, we continued to use fresh media with glucose to prepare our standards.

Finally, we checked if spiking with one cofactor impacted measured concentrations of the other cofactors, which could indicate interconversion between cofactors. We tested this with two high concentrations (1 or 5 μM). This revealed only minimal or no effects on other cofactor concentrations in most cases (Figure S3). Exceptions were NADPH, which resulted in increased NADP^+^ levels: app. 5% of NADPH seems to be oxidized to NADP^+^ under the used reaction conditions, which led to a 48% or 16% increase in measured NADP levels when 5 μM or 1 μM NADPH was spiked, respectively. This was only observed when high concentrations of the cofactors were spiked, however, typically between 0.1 to 0.5 μM NADPH and NADP^+^ are measured. Surprisingly, spiking NMN and NADP^+^ had a consistent negative effect on NAD^+^ concentrations. NAD^+^ concentrations decreased with 23 - 24%, when 5 μM of NMN or NADP^+^ was spiked. When 1 μM NMN or NADP^+^ was spiked, the observed decrease in NAD^+^ was only 10%. Also, minor fractions (<1%) of NAD^+^ and NADP^+^ were degraded to NMN. Based on the combination of these small fractions and the typical concentrations measured in biological samples, these degradation effects fall within the detection limits of our method.

### Comparison of the HPLC-FLD method with INT assay

To benchmark our method, we grew *E. coli* on different carbon sources (glucose, glycerol, and acetate) and measured the cofactor levels. Since colorimetric enzyme assays are a commonly used to measure NAD^+^ and NADH levels ^50^, we used the INT method for comparison ^25^ (Figure 3).

**Figure 3.**
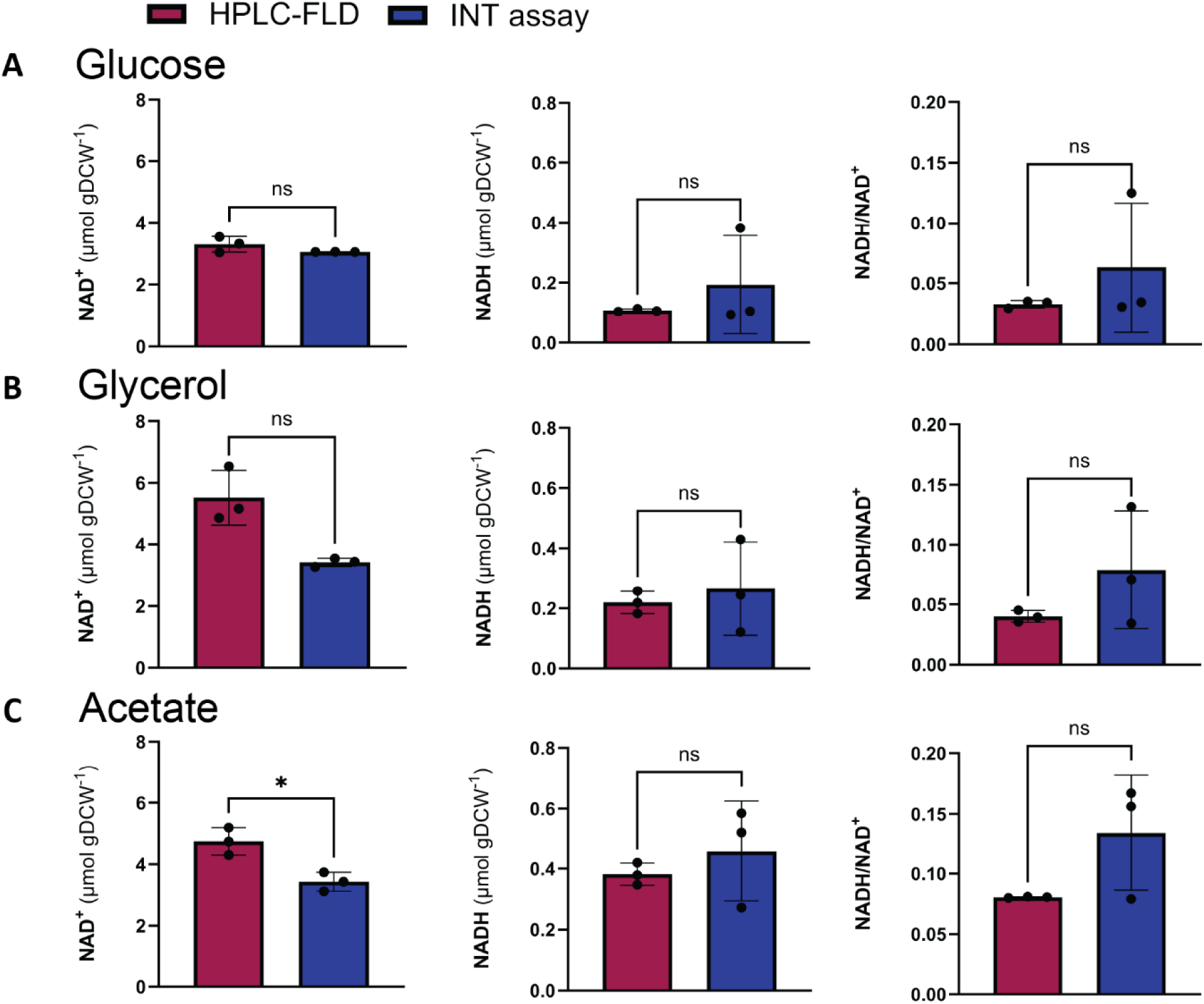
Comparison of cofactor levels and ratios measured in *E. coli* during exponential growth on different carbon sources using two methods. A) Cultures grown on 20 mM glucose, B) cultures grown on 40 mM glycerol, C) cultures grown on 60 mM acetate. In red, the values measured with the newly developed HPLC-FLD method and in blue the values measured using the INT assay. Each dot represents the average of a biological replicate measured in two (HPLC-FLD) or six (INT-assay) technical replicates. Bars show the average of the biological triplicates with their corresponding standard deviation. Significant differences, indicated with *, between the two methods were determined with a Welch’s t-test (p < 0.05).

Most values obtained for cellular cofactor concentrations for the INT were in the same range as the concentrations measured with HPLC-FLD and did not differ significantly. The measured NAD^+^ concentrations for glucose-grown *E. coli* differed by 10%, while for glycerol and acetate, the differences between the NAD^+^ levels measured with INT assay and HPLC-FLD were larger between 30 – 40%. Only for acetate-grown these differences were significant (Figure 3C). Overall, the HPLC-FLD method shows comparably but slightly higher NAD^+^ values than the INT assay.

NADH levels were harder to measure accurately with the INT assay, resulting in larger standard deviations. The standard error for the HPLC-FLD measurements were 0.03-0.022, while for the INT assay the standard error were 0.090-0.096. Therefore, we also tested the LOQ for the INT assay. As expected, the NAD^+^ levels measured in the *E. coli* grown cells on the different carbon sources fell in the range of the determined LOQ for the INT assay (Table S3). The LOQ for NADH was higher than the levels measured in *E. coli* cultures, meaning that NADH levels could not be reliably determined with the INT assay.

The NADPH and NADP^+^ levels were not determined using an INT assay. Results from the HPLC-FLD method showed that NADPH/NADP^+^ ratios were similar in glucose- and acetate-grown cells (0.57 ± 0.1 and 0.8 ± 0.1, resp), but 2-3-fold higher in cells grown on glycerol (1.8 ± 0.3) (Figure S5).

### Application example: cofactor levels in engineered NMN-producing strains

As an application, we showed that our method can be used to measure cofactor levels and NMN in genetically engineered *E. coli* strains. Nicotinamide mononucleotide (NMN) is an intermediate in the NAD-biosynthesis pathways and has health-beneficial properties against age-related diseases and diabetes ^51,52^. In addition, NMN can be used as a non-canonical redox cofactor ^53–56^. For these reasons, NMN has been overproduced in *E. coli* ^35–37,57–59^. Here, we analyzed NAD(H), NADP(H) and NMN levels in a strain overexpressing the NMN-synthetase NadV from *Francisella tularensis* (FtNadV) (Figure 4A). To reduce NMN degradation, the NMN amidohydrolase PncC was deleted in this strain ^57^. The resulting *E. coli* Δ*pncC* FtNadV strain was grown on minimal media with glucose with or without 1 mM nicotinamide (NAM) to assess the effect on the intracellular NAD(P)(H) and NMN levels.

**Figure 4.**
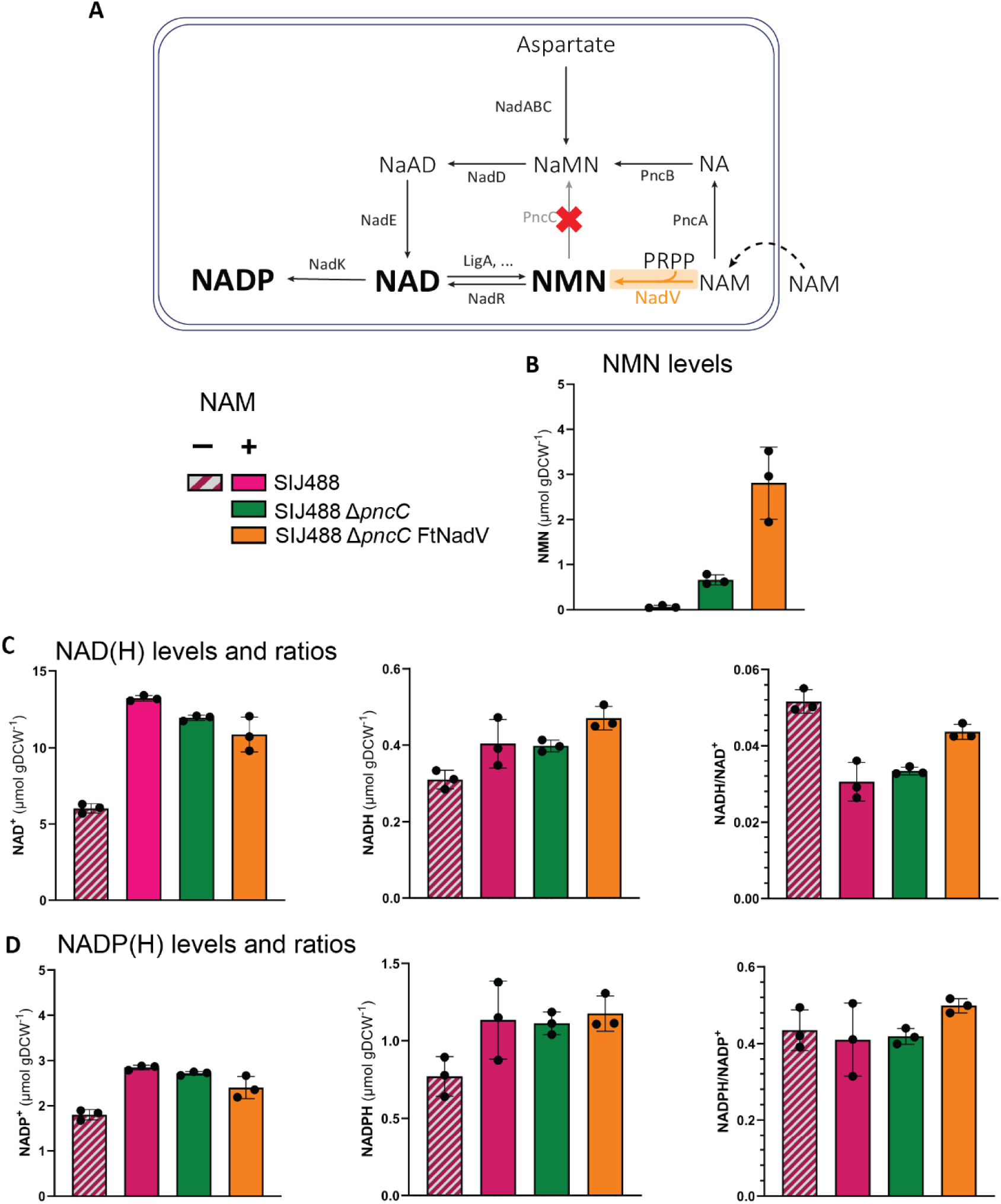
Cofactor levels in NMN overproducing mutant strains grown on M9 medium with 20 mM glucose, with or without 1 mM nicotinamide supplied. A) metabolic scheme of mutations in mutant strains. B) NMN levels in strains. C) NAD(H) levels and ratios measured in strains. D) NADP(H) levels and ratios measured in strains. Each data point represents an average of technical duplicates. The average of the biological triplicates is represented in bars with their corresponding standard deviation.

The strain overexpressing FtNadV accumulated NMN to 2.2 ± 0.63 μmol gDCW^-1^ in the presence of NAM (Figure 4B). This intracellular level corresponded to half the concentration of NAD^+^ in the reference strain without NAM added (Figure 4C). Notably, NAD^+^, NADP^+^ and NADPH increased 1.5- to 2-fold upon feeding NAM in all three strains (Figure 4C, D). NADH levels also increased, however not as much as the other cofactors, resulting in a lower NADH/NAD^+^ ratio. This shows that our method can be used to study the effects of engineering strategies on individual cofactor levels and ratios and NMN levels, including in NMN-overproducing strains.

### Cross laboratory and species validation: cofactor levels in *P. putida*

To further demonstrate the applicability of the developed HPLC-FLD method, we used it to analyze NAD^+^ and NMN levels in *Pseudomonas putida*, an alternative microbial chassis. Furthermore, these analyses were done in a different laboratory to also illustrate ease of implementation. We selected *P. putida*, because this soil bacterium is endowed with a highly versatile enzymatic repertoire and notable metabolic robustness, which make it an attractive platform for industrial biotechnology ^60^. Although the NAD⁺ biosynthetic pathway in *P. putida* has not been fully characterized, we transplanted the same strategy to this organism to increase NMN levels. For this, the nicotinamide-nucleotide amidohydrolase (*pncC*) encoding gene was deleted and nicotinamide phosphoribosyltransferase (*nadV*) was expressed under the control of a rhamnose-inducible promoter (pP4S·*nadV*).

Comparative analysis of intracellular NAD-derived metabolites in the parental and engineered strains revealed trends similar to those observed in *E. coli*. Intracellular NMN concentrations increased from nearly undetectable levels to approximately 2.6 ± 0.5 µmol gCDW⁻¹ (Figure 5A). In addition, the intracellular concentration of NAD^+^ rose from approximately 4.1 ± 0.7 to 7.5 ± 1.2 µmol gCDW^-1^ (Figure 5B) . Furthermore, we could observe that the intracellular nicotinamide-based redox cofactor pools are slightly lower than in *E. coli,* which could be explained by its central metabolism being highly dependent on PQQ-dependent oxidoreductases ^61^. Importantly, we demonstrated that intracellular concentrations determined using the INT and HPLC-FLD methods were also comparable in this microorganism without the need of method readaptations (Figure 5C).

**Figure 5.**
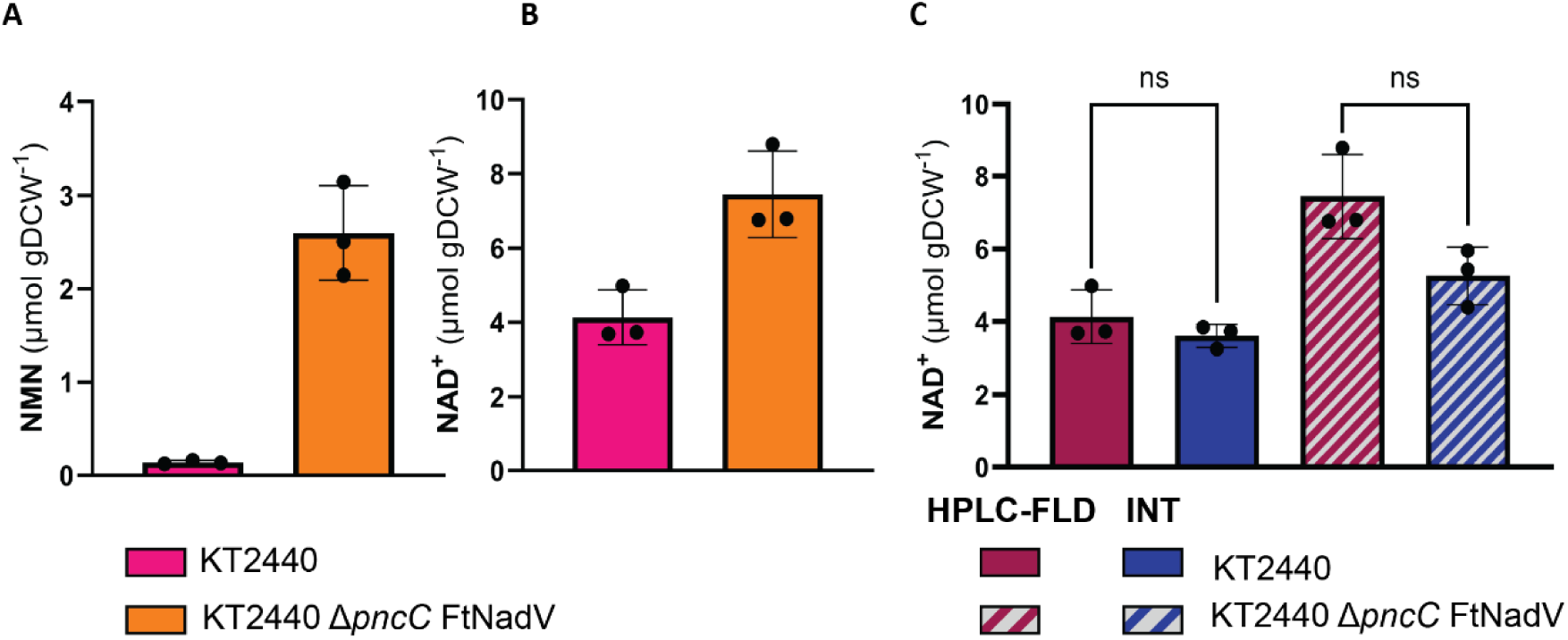
NMN and NAD^+^ levels measured in *P. putida* strains. A) NMN levels measured HPLC-FLD. B) NAD^+^ levels measured with HPLC-FLD. C) NAD^+^ levels measured with HPLC-FLD and INT-assay in *P. putida* strains. The average of the biological triplicates is represented in bars with their corresponding standard deviation. Significant differences, indicated with *, between the two methods were determined with a Welch’s t-test (p < 0.05).

## Discussion

We developed a quantification method for NAD(P)(H) and NMN levels and ratios based on HPLC and fluorescence detectors. The method was thoroughly characterized using *E. coli* cultures, and this method can be used to characterize nicotinamide redox cofactor levels in different physiological conditions and mutant strains.

Extensive characterization of the method showed that LOQs for this method (< 20 nM, Table S1) are close to LOQs reported in papers describing LC/MS based protocols (11.5 – 173 nM ^62,63^). In addition, spiking *E. coli* samples proved that this method can be used to accurately measure NAD^+^, NADP^+^, and NMN. For NADH and NADPH, we observed a reduced recovery of the spiked concentrations. However, within the range tested, concentrations scaled linearly (Pearson correlation r > 0.996, p-value < 0.0001). Of note is that we observed that the matrix, e.g., the medium conditions, can have a strong influence on measured fluorescence levels. This highlights the importance of including correct controls to assess the influence of the matrix on fluorescence intensity, which is already common practice in the development of many LC/MS approaches ^29,64,65^.

To benchmark this method and show that this method can be applied to measure cofactor levels in *E. coli* under different conditions, we measured NAD^+^ and NADH in cells grown on different carbon sources with the INT assay and HPLC-FLD-based method (Figure 3). The HPLC-FLD method gave similar results as the INT assay for NAD^+^ and NADH levels for most of the tested conditions. Between carbon sources, little differences were seen regarding the different cofactor levels. Also, in a different laboratory and bacterium, *P. putida*, both methods gave similar results (Figure 5C). The HPLC-FLD method, however, outperformed the INT assay in different aspects. First, CoVs were similar for NAD^+^ (on average 11.1 ± 4.5% and 4.4 ± 4.5%, p-value 0.14) or significantly lower for NADH (on average 10.5 ± 6.3% versus 60 ± 24.4%, p-value 0.027) compared to the INT assay. In addition, the LOQs for the HPLC-FLD method are lower than for the INT assay.

The measured NADH/NAD^+^ ratios in *E. coli* grown on different carbon sources were in the range of previously reported values (0.007 – 0.05) ^2,66^. However, reported NADPH/NADP^+^ ratios in these previous reports were far above 1 (10 to 50), while we detected ratios between 0.5-1.7. This difference can mostly be attributed to around 20-fold differences in reported NADP^+^ levels. It is commonly known that NAD(H) is predominantly present in the oxidized form, while NADP(H) is predominantly present in the reduced form in cell metabolism ^3,67^. However, NADPH/NADP^+^ ratios around or below 1 in *E. coli* have also been reported _62,63,68–70._

Furthermore, we showed that the method is especially suitable to measure NMN and NAD(P)(H) in mutant strains where larger differences are expected between cofactor levels. NMN levels measured in our Δ*pncC* FtNadV mutant strains were similar to values reported in literature ^37,57^. In addition, we showed that NAD^+^ and NADP^+^ levels doubled upon feeding NAM. Fluorescent derivatization of NMN has been used to measure cellular NMN concentrations with plate reader. However, using plate readers it is not possible to distinguish between NAD^+^, NADP^+^, and NMN fluorescence ^35,36^. While literature assumes that NAD^+^ and NADP^+^ are highly regulated, and levels do not change in engineered strains ^37^, our data clearly indicate otherwise: cellular NAD^+^ and NADP^+^ levels did increase when nicotinamide was present in the medium. Therefore, measuring fluorescent derivatized NMN using plate readers should be done with caution, and is only suitable for extracellular NMN levels.

While our data clearly indicates that NAD^+^, NADP^+^, and NMN can be precisely quantified, results for NADH and NADPH are less precise. More degradation of NADH and NADPH in biological samples was observed compared to standards, which has been noted before ^49^. Alkaline extraction of these reduced cofactors is commonly used in enzymatic assays, and to completely degrade NAD^+^ and NADP^+^, the samples need to be heated. Here, this was performed at 100 °C, which presumably leads to a higher instability of these cofactors ^71^.

Despite its limitations, this method is an innovative and rapid addition to already existing methods to measure cofactor levels and ratios in bacterial cultures. Redox cofactors remain challenging to measure, due to their quick turnover, low concentrations, and instability. This HPLC-FLD protocol gives a more complete view of cofactor levels and ratios than the classic INT assay, since it can measure NAD^+^, NADP^+^, NADH, NADPH, and NMN in the same sample. The method has the potential to be further developed to other NAD^+^ metabolites and non-canonical redox cofactors, such as NR, NMNH, NCD(H). In addition, the equipment needed is more affordable, and the data is easier to analyze than LC/MS-based methods. Lastly, the derivatization and detection methods are cell agnostic and can be translated to other cell types, e.g., mammalian cells. To conclude, our method is especially valuable to measure nicotinamide cofactor levels in many samples in a fast and reliable comparative manner, as is commonly needed in fields like metabolic engineering.

## Acknowledgements

The authors would like to thank Wendy Evers for technical support and Matic Kostanjsek, Jenny Bakker and Rutger Verbakel for valuable discussions.

