## Supplementary information for "A rapid HPLC-based method to determine NAD(P)(H) and NMN redox cofactor concentrations and ratios in microbes"

### FtNadV - sequence

ATGAGCTTTGATAATCTATTATTAATGACCGATAGCTATAAGCATAGCCATAGGTATCAGTATCCGCGCGATACCCATTA
TTTACATTTTTATTTAGAAAGCCGGGGCACCGCCAACAAAGATTTAGGCAACTATACCAAATTTTTTGGCTTACAGTATTA
TGTTAAAAAATATCTGAGCCAGCCGATCACCCAGCAGATGATCGATGATGCCGAAAAAATTCTGCTGGCGCATGGCCTG
CCGTTTTATCGGAGCGGGTTTGAGAAGATCCTGAACAACTACAACGGATATCTGCCGATTCGCATTCGGGCCGTGCGC
GAGGGCAGCTTAATTCCGCTGCATAACGTGTTAATGACCATCGAAAGCACCGATGAAGAATTATTTTGGCTGCCGGGGT
TTGTGGAAACCCTGTTACTGAAAGTTTGGTATCCGACCACCGTTGCCACCATTAGCTTTAACATCAAACAATTAATCAAG
CGCTATTTATTAGAAACCGCCGATAGCTTAGATAAACTGGATTTTATGTTACATGATTTCGGCTATCGCGGCGTGTCAAG
CGAAGAGAGCGCGGGAATTGGCGGAGCTGCCCACTTAACCAACTTCTTAGGGACCGATACCCTGGCCGCCCTGCATGT
GTGTAAAGAATTCTATGCGGAAGATATGGCGGGGTTTAGCATACCGGCCAGCGAGCATAGCACCATGACCTCATGGGG
CGTAGGAACCGAGTGCGAGCGGGAGGCGTTTGAAAACATGATCGCCCAGTTTGGCGATAGCAGCGTGTTATATGCGTG
CGTGAGCGATAGCTGGGATTTTAAAAAAGCCATTCAGACCTGGGTGGATCTGAAAGATCGAGTGACCGCGAAAAAAGC
CAACCTGGTGATACGACCGGATAGCGGGGATGCTGTGGATAACATATTATATGCCTTATATGAACTGGATAAAGGCTAT
GGCAGCCGCTTAAATAGCAAAGGGTATAAAGTTCTGAATAACGTGGCCTTAATTCAGGGCGATTCAGTTAGCATTTCAT
TAGCCAAAAAAGTACTGGAAGCGATGAAAATCCAGGGCTACAGCGCCGAGAACATAGCGTTTGGCATGGGCGGAGCTC
TGCTACAGGGCAACTATGAATCAAGCATCAACCGCGATAGCTTCAAATTTGCCATAAAGTGTAGCGCTATTATGCGGGG
CAACACCTTAATCGGCGTAAAAAAAGAACCGATTACCGACTTAGCCAAAAAAAGCAAACAGGGGCGATTAGATCTGATC
AAAGATGCGAAGGGAAACTATAAAACCATCGTGTTAGATGATAGCTATGCGTTAGGAGAATATCATCCGGAGAGCCAGT
TACAGACCTATTATGATAACGGCGAAATTAAATTTGAGCAAAGCCTGGCCCAAATCCGGAACTACACCAACTAA

Table S1. Strains and plasmids used.

| Strain | Characteristics | Reference |
|---|---|---|
| ***E. coli* DH5α λpir (Cloning Host)** | DH5α λpir | New England Biolabs |
| ***E. coli* JW2670** | *E. coli* K-12 BW25113 Δ*pncC*::FRT_Km_FRT | [38] |
| ***E. coli* SIJ488** | K-12 MG1655 Tn7::para-exo-beta-gam; phra-FLP; xylSpm-IsceI | [39] |
| ***E. coli* SIJ488 Δ*pncC*** | SIJ488 Δ*pncC::FRT* | This study |
| ***E. coli* SIJ488 Δ*pncC* FtNadV** | SIJ488 Δ*pncC::FRT* pSevab32-FtNadV | This study |
| ***P. putida* KT2440** | KT2440 | This study |
| ***P. putida* KT2440 Δ*pncC* FtNadV** | KT2440 Δ*pncC* pP4S·*nadV* | This study |
| **Plasmids** | | |
| **pSb1C3 (standard IGEM Biobrick vector)** | Cm, ColE1 | https://parts.igem.org/Help:2019_DNA_Distribution |

| | | |
|---|---|---|
| **pSEVAb32** | Cm, RK2 | [41] |
| **pSEVAb32-FtNadV** | Cm, RK2, J23100-Bbab0034-FtNadV | This study |
| **pSNW2** | Suicide vector for deletions in Gram-negative bacteria, Km, oriT, traJ, lacZα, ori(R6K), P14g-BCD2-msfGFP | [42] |
| **pSNW2·*pncC*** | Derivative of pSNW2, carrying HRs to delete *pncC* | This study |
| **pQURE6H** | Conditionally replicating vector; derivative of pJBSD1, Gm, XylS/$P_m$-ISceI and P14g-BCD2-mRFP | [42] |
| **pP4S·*nadV*** | Sm, pBBR1, RhaSR/PrhaBAD- FtNadV | This study |

Table S2. Primers used. Capital letters indicate BsaI recognition sites.

| **Name** | **Sequence** |
|---|---|
| **OP1_BBaB0034_FtNadV** | cttGGTCTCaactagagaaagaggagaaatactagatgagctttgataatc |
| **SEVA_FtNadV_Rv** | aGGTCTCgactgcagcggccgctactagtattattagttggtgtagttccg |
| **SEVA_EvBBa_J23100** | aGGTCTCctagtagctagcactgtacctaggactgagctagccgtcaactctagaagcgg |
| **EV** | ccgGGTCTCaattccagaaatcatccttagcgaaagc |
| **PV** | aGGTCTCacagtccggcaaaaaaggg |
| **KO_PncC_R** | gctgggcaaccatcaacaag |
| **KO_PncC_F** | gataacgtctactgcgccagaac |

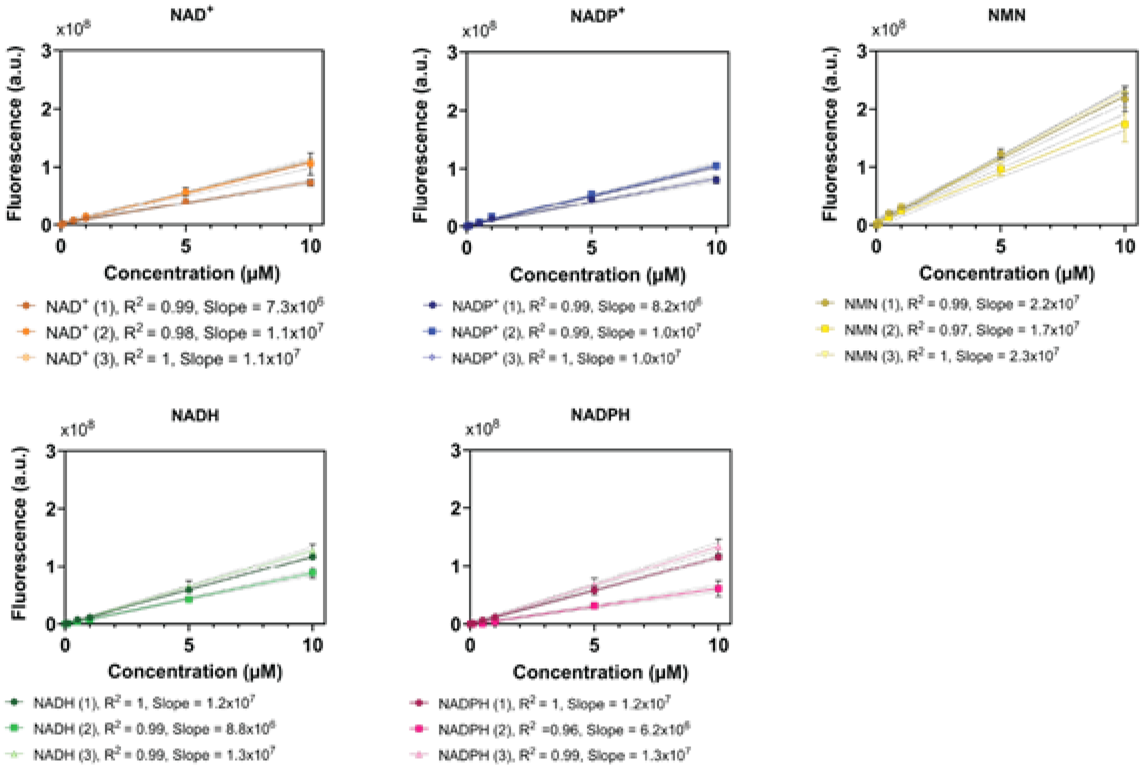


Figure S1. Standard curves of the different peaks from standards diluted in M9 with 20 mM glucose. Standard were prepared three times in duplicate and run during different runs. The average is shown of the duplicates with corresponding standard deviation.

Table S3. LOD and LOQ in µM measured in standards prepared in M9 with 20 mM glucose. For three standard curves LOD and LOQ were determined. Highest LOD and LOQ found over the three different runs are shown here.

| | *LOD (3.3= S/N)* | *LOQ (10 = S/N)* |
|---|---|---|
| **$NAD^+$** | 0.004 | 0.011 |
| **$NADP^+$** | 0.005 | 0.015 |
| **NMN** | 0.002 | 0.006 |
| **NADH** | 0.004 | 0.013 |
| **NADPH** | 0.006 | 0.018 |

Table S4. Average recovery for each concentration of compound spiked. Recovery was calculated by: $R = (A_{spiked} - A_{unspiked}) / A_{standard} * 100\%$. Each concentration was measured in triplicate. Average recovery is shown with coefficient of variation (CoV%).

| | Concentration spiked (µM) | Average Recovery (%) | CoV (%) |
|---|---|---|---|
| **$NAD^+$** | 0.1 | 69.8 | 52.00 |
| | 0.5 | 105.3 | 3.68 |
| | 1 | 106.9 | 3.67 |
| | 5 | 93.0 | 3.47 |
| **$NADP^+$** | 0.1 | 81.4 | 32.77 |
| | 0.5 | 102.0 | 0.72 |
| | 1 | 96.0 | 3.99 |
| | 5 | 100.4 | 8.33 |
| **NMN** | 0.1 | 94.2 | 4.16 |
| | 0.5 | 95.1 | 3.59 |
| | 1 | 114.3 | 14.15 |
| | 5 | 125.9 | 5.83 |
| **NADH** | 0.1 | 66.5 | 44.35 |
| | 0.5 | 77.7 | 21.81 |
| | 1 | 63.6 | 16.19 |
| | 5 | 77.4 | 7.90 |
| **NADPH** | 0.1 | 79.1 | 3.87 |
| | 0.5 | 74.4 | 5.38 |
| | 1 | 65.6 | 10.95 |
| | 5 | 81.7 | 1.36 |

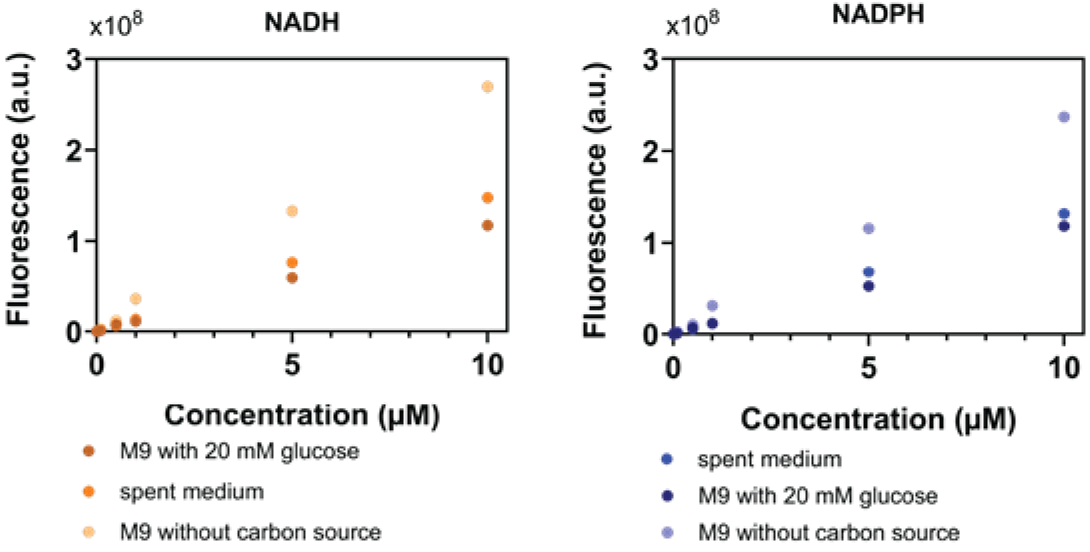


Figure S2. Influence of glucose on standard curves of the reduced cofactors. Spent medium was retrieved from filtered exponentially growing *E. coli* cultures.

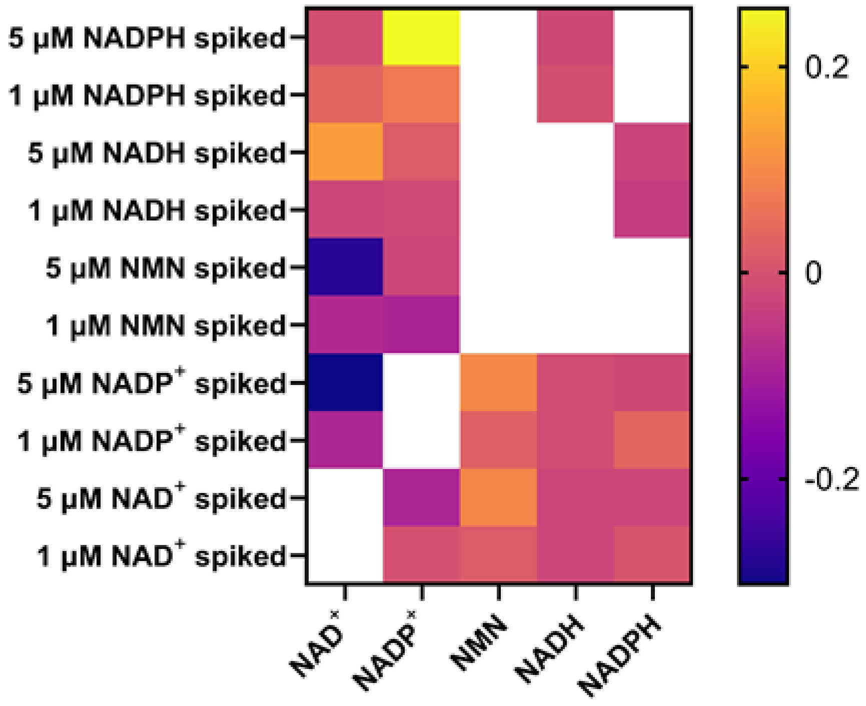


Figure S3. Average change in concentration (μM) of cofactors upon spiking of NAD(P)(H) and NMN. The tiles represent the average of triplicate measurements.

Table S5: Limit of detection and limit of quantification calculated for measurements of cells grown on the three different carbon sources glucose, glycerol and acetate. Values are given as averages (± standard deviation) of three biological replicates with $OD_{600}$ 0.7.

| | **LOD $NAD^+$ (µmol $gDCW^{-1}$)** | **LOQ $NAD^+$ (µmol $gDCW^{-1}$)** | **LOD NADH (µmol $gDCW^{-1}$)** | **LOQ NADH (µmol $gDCW^{-1}$)** |
|---|---|---|---|---|
| **Glucose** | 0.63 (± 0.12) | 1.92 (± 0.35) | 2.00 (± 0.31) | 6.05 (± 0.93) |
| **Glycerol** | 0.96 (± 0.68) | 2.92 (± 2.05) | 0.81 (± 0.44) | 2.46 (± 1.32) |
| **Acetate** | 1.31 (± 0.42) | 3.96 (± 1.29) | 0.82 (± 0.53) | 2.50 (± 1.61) |

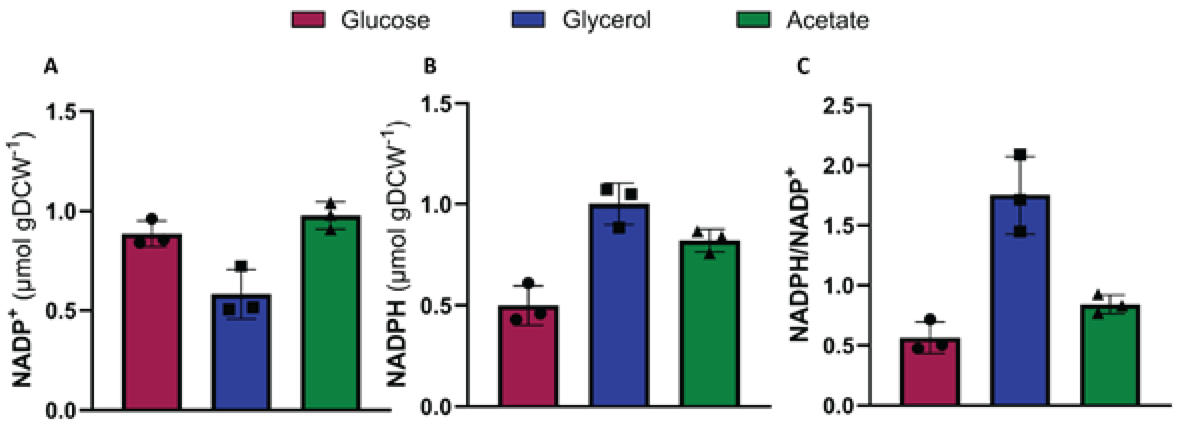


Figure S4. NADP(H) levels and ratios for *E. coli* cultures grown on different carbon sources measured with HPLC-FLD: $NADP^+$ (A), NADPH (B) and NADPH/$NADP^+$ (C). Experiments were performed using samples from biological triplicates measured in duplicates. The data points correspond to the averages of technical duplicates. The bars correspond to biological triplicates with corresponding standard deviation.
